# Long-term stability of avalanche scaling and integrative network organization in prefrontal and premotor cortex

**DOI:** 10.1101/2020.11.17.386615

**Authors:** Stephanie R. Miller, Shan Yu, Sinisa Pajevic, Dietmar Plenz

**Affiliations:** Section on Critical Brain Dynamics, National Institute of Mental Health, Bethesda, MD, USA; Brainnetome Center, Institute of Automation, Chinese Academy of Sciences, China; Section on Quantitative Imaging and Tissue Sciences, National Institute of Child Health and Development, NIH, Bethesda, MD, USA

**Keywords:** nonhuman primate, resting activity, criticality, brain dynamics, neuronal avalanches, integrative network organization

## Abstract

Ongoing neuronal activity in the cortex establishes functional networks of synchronization that reflect normal and pathological brain function. The reconstruction of these networks typically suffers from the use of indirect measures of neuronal activity at low spatiotemporal resolution and a lack of longitudinal tracking. Accordingly, the precise nature of the underlying synchronization dynamics and its translation into robust graph theoretical markers are not well characterized. Here, we studied the stability of cortical dynamics and reconstructed functional networks over many weeks in prefrontal and premotor cortex of awake nonhuman primates. We monitored neuronal population activity directly in the ongoing local field potential (LFP) at high spatial and temporal resolution using chronically implanted high-density microelectrode arrays. Ongoing activity was composed of neuronal avalanches exhibiting stable, inverted parabolic profiles with the collapse exponent of 2 in line with a critical branching process. Avalanche-based functional networks, reconstructed using a Normalized Count estimator, revealed robust integrative properties characterized by high neighborhood overlap between strongly connected nodes and robustness to weak-link pruning. “Entropy of mixing” analysis demonstrated progressive link reorganization over weeks. The long-term stability of avalanche scaling and integrative network organization should support the development of robust biomarkers to characterize normal and abnormal brain function.

## Introduction

Ongoing neuronal activity in the mammalian brain gives rise to distinct functional networks that reflect normal and pathological brain function (Bullmore and Sporns, 2009; Fox and Raichle, 2007; Raichle, 2015). Empirical measures of these networks, however, are often limited in the precision with which they reflect the organization of neuronal population activity and a lack of longitudinal follow-up. Networks are either reconstructed from indirect non-neuronal activity at low temporal resolution using functional magnetic resonance imaging (fMRI), or via more direct neuronal activity measures such as magnetoencephalography (MEG) and the electroencephalogram (EEG) which suffer predominantly from low spatial resolution (Bassett and Sporns, 2017). Accordingly, there is limited knowledge about the stability and nature of the spatiotemporal dynamics and corresponding functional connectivity of local neuronal populations that give rise to fluctuations in ongoing cortical activity.

In contrast to macroscopic and indirect methods, neuronal activity measured at single-cell resolution could in principle allow for precise and robust spatiotemporal reconstruction of functional networks in cortex. For example, in macaque cortex single-unit activity can be recorded over many weeks (Nicolelis et al., 2003) and identifies firing statistics of neurons over days (Dickey et al., 2009; Jackson and Fetz, 2007). Advanced approaches may involve tetrodes, automated classifiers (Tolias et al., 2007) or second order statistics as classifiers (Fraser and Schwartz, 2011). Even advanced single-unit approaches, however, limit robust tracking of activity to small groups of individual neurons. Chronic expression of genetically-encoded calcium indicators (GECIs) can increase the neuronal count and identify orientation-selective neurons over months (Li et al., 2017), yet at low temporal resolution. The local field potential (LFP), alternatively, provides access to an intermediate spatial resolution of cortical activity that is between the macroscopic scale of fMRI, MEG and EEG and single cell accuracy. It robustly measures local neuronal population activity with high temporal and reasonable spatial resolution of several hundred micrometers (e.g. (Bellay et al., 2021; Katzner et al., 2009)) over many weeks. In nonhuman primates (NHPs), the LFP has been shown to correlate well with local unit-activity (Donoghue et al., 1998; Petermann et al., 2009; Rasch et al., 2008) and offers robust brain-machine interface performance (Mehring et al., 2003).

It is now well established that the ongoing LFP in superficial layers of nonhuman primates organizes into spatiotemporal activity clusters whose sizes and durations distribute according to power laws (Miller et al., 2019; Shew et al., 2009; Yu et al., 2017), the hallmark of neuronal avalanches (Beggs and Plenz, 2003). The spatiotemporal correlations underlying neuronal avalanches give rise to functional “integrative” networks, in which the topology and weights of links organize such that end-nodes of strong links are more likely to have many common neighbors, as quantified by a high correlation between the link weight and the link clustering coefficient ((Pajevic and Plenz, 2012), *cf*. edge-clustering coefficient in (Radicchi et al., 2004)), while being robust to the loss of weak links, in terms of maintaining the advantages of a small-world topology (Watts and Strogatz, 1998). This specific arrangement describes many complex networks ranging from genetic networks to social, i.e. “friendship”, networks (Pajevic and Plenz, 2012). Importantly, integrative networks can form using local plasticity rules that operate on neuronal avalanches (Alstott et al., 2015) in which weak links provide a large basin for plastic changes as they can be removed or rewired, permitting flexibility, while preserving the small-world network property. The stability over days and weeks of cortical avalanche dynamics and corresponding integrative networks has not been explored for cortex. Size and duration distributions of avalanches do not consider the spatiotemporal organization of avalanches, thus multiple functional connectivities could give rise to similar power law statistics. Likewise, integrative networks allow for the reorganization of individual links while maintaining global integrative properties (Pajevic and Plenz, 2012).

Here we recorded ongoing LFP activity in prefrontal and premotor cortex of NHPs over weeks using chronically implanted high-density microelectrode arrays. We demonstrate stable avalanche dynamics with power law statistics, the temporal avalanche profile of an inverted parabola and corresponding scaling exponent of 2, which is the expected value for critical branching process dynamics (di Santo et al., 2017; Miller et al., 2019; Sethna et al., 2001). We further demonstrate that these avalanches give rise to non-trivial integrative functional networks despite progressive reorganization of the link strengths as identified using Normalized Count (NC) estimators (Pajevic and Plenz, 2009) and the “entropy of mixing” analysis (see Material and Methods). Our results suggest that avalanche dynamics and corresponding integrative network organization identify a robust cortical state in the adult brain, which should inform dynamical models of brain function and might allow for the early identification of pathological brain states.

## Results

The spatiotemporal organization of the ongoing LFP (1 – 100 Hz) was monitored with chronically implanted high-density microelectrode arrays (~10 × 10 electrodes; interelectrode distance 400 μm) in premotor (PM, n = 2 arrays) and prefrontal cortex (PF, n = 4 arrays) of three macaque monkeys (*Macaca mulatta*; K, V, N; Fig. 1A, B). The animals were awake and seated in a monkey chair during the recording sessions but were not given any specific stimulus or task. On average about 4 ± 2 hr of LFP activity during 9 ± 7 recording sessions over the course of 5 ± 4 weeks were analyzed per monkey (85 ± 8 electrodes/array; Fig. 1C).

**Figure 1:**
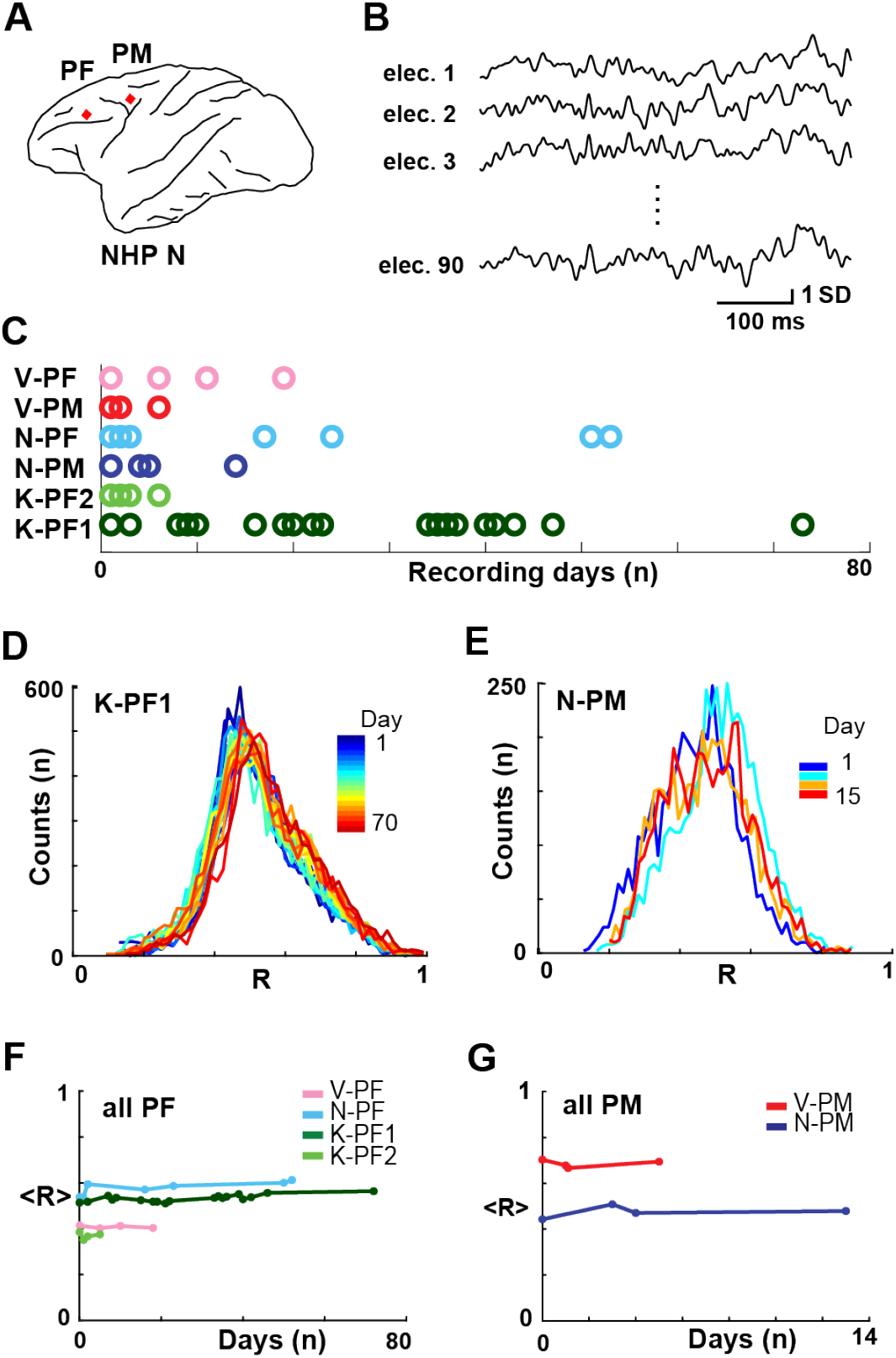
Ongoing cortical dynamics are stable over weeks in superficial layers assessed by pairwise correlations in the LFP. **A,** Sketch of chronic prefrontal (*PF*) and premotor (*PM*) cortical implantation sites (*red diamonds*) for high-density microelectrode arrays. **B,** Example of simultaneously recorded ongoing LFP (1 – 100 Hz) at single electrodes with correlated activity. **C,** Overview of ~30 min recording sessions per day (*color coded circles*) over several weeks for each NHP. **D, E,** Pair-wise correlations of the continuous LFP distribute similar over time. Normalized histogram of pairwise correlations between all functioning channels for each recording day (*color bar*) in two NHPs (*left*: prefrontal recording in K; *right*: premotor recordings in N). **F, G**, The average correlation, <*R*> (from *D*, *E*), was constant over weeks in PF (*left*) and PM (*right*) arrays.

### Stability of average pairwise correlation and avalanche dynamics in prefrontal and premotor cortex

The pairwise Pearson’s correlation, *R*, is a standard measure to quantify the correlation in neuronal activity between distant cortical sites. We found that for the continuous LFP, the distribution of *R*, obtained from ~5,000 electrode comparisons per array, was broad yet similar across days (Fig. 1D, E). The average correlation, <*R*>, while different between arrays, was constant over many days and weeks (Fig. 1F, G; linear regression fit of 0.4 ± 0.7 ×10^−2^ for all arrays; see Table 1 for individual arrays).

**Table 1.** Summary of average estimates for the mean Pearson’s pairwise correlation coefficient, <*R*>, the exponent α of avalanche size distributions for each nonhuman primate and array at temporal resolutions of *Δt* = 1 ms and *Δt* = 30 ms respectively. Exponents were derived from maximum likelihood estimates. Stability of α over all recording days was quantified by the linear regression slope within 95% confidence limits (CfL).

| | $\langle R \rangle$<br>Stability | Avalanche<br>size<br>distribution<br>($\Delta t = 1$ ms) | slope $\alpha$<br>stability<br>($\Delta t = 1$ ms) | Avalanche<br>size<br>distribution<br>($\Delta t = \langle IEI \rangle$) | slope $\alpha$<br>stability<br>($\Delta t = \langle IEI \rangle$) | Avalanche<br>size<br>distribution<br>($\Delta t = 30$ ms) | slope $\alpha$<br>stability<br>($\Delta t = 30$ ms) |
| --- | --- | --- | --- | --- | --- | --- | --- |
| | Regression<br>slope (95%<br>CfL) ($\times 10^{-3}$) | Exponent $\alpha$<br>(± SD) | Regression<br>slope (95% CfL)<br>($\times 10^{-3}$) | Exponent $\alpha$<br>(± SD) | Regression<br>slope (95%<br>CfL) ($\times 10^{-3}$) | Exponent $\alpha$<br>(± SD) | Regression<br>slope (95%<br>CfL) ($\times 10^{-3}$) |
| **V-PF** | -1 (-3, 2) | $1.71 \pm 0.00$ | 0 (-1, 0) | $1.26 \pm 0.01$ | -2 (-3, 3) | $0.65 \pm 0.03$ | -3 (-9, 2) |
| **V-PM** | 0 (-62, 62) | $1.64 \pm 0.02$ | 6 (-40, 51) | $1.29 \pm 0.04$ | 16 (-28, 60) | $1.12 \pm 0.05$ | 20 (-2, 43) |
| **N-PF** | 1 (0, 2) | $1.71 \pm 0.02$ | -1 (-1, 0) | $1.41 \pm 0.01$ | 0 (-1, 0) | $0.97 \pm 0.02$ | 0<br>(-1, 1) |
| **N-PM** | 1 (-13, 15) | $1.73 \pm 0.02$ | 2 (-8, 12) | $1.31 \pm 0.02$ | 0 (-1, 1) | $0.75 \pm 0.02$ | 3 (-6, 12) |
| **K-PF1** | 1 (0, 1) | $1.65 \pm 0.02$ | -1 (-1, 0) | $1.29 \pm 0.01$ | 0 (-1, 0) | $0.90 \pm 0.02$ | 0 (0, 1) |
| **K-PF2** | 0 (-20, 20) | $1.77 \pm 0.03$ | 5 (-33, 43) | $1.36 \pm 0.02$ | 1 (-27, 29) | $0.74 \pm 0.03$ | 10 (-16, 35) |
| **Mean ±**<br>**SD** | **$0 \pm 1$** | **$1.70 \pm 0.05$** | **$2 \pm 3$** | **$1.32 \pm 0.05$** | **$3 \pm 7$** | **$0.86 \pm 0.17$** | **$5 \pm 9$** |

We next explored the stability of neuronal avalanche dynamics by discretizing the LFP from ongoing activity in the NHPs (Petermann et al., 2009; Yu et al., 2017). We previously demonstrated that negative deflections in the continuous LFP signal (nLFPs) track the local neuronal population activity and correlate with extracellular unit firing (Bellay et al., 2021; Petermann et al., 2009). Accordingly, discrete nLFPs were defined by crossing a threshold of −2 SD and extracting their peak times (Fig. 2A). nLFPs were then grouped into avalanches at temporal resolution *Δt* of the mean time interval of successive nLFPs on the full array, <IEI>, which ranged between 3 – 4 ms (Fig. 2B; see Material and Methods). We found that the probability density distribution of avalanche size, which typically ranges from size 1 to the number of electrodes for each array (Yu et al., 2011), exhibited power laws for each day of recording (Fig. 2C, D; *Δt* = <IEI>, truncated power law vs. exponential, LLR >> 1, *p* < 0.005; or vs. log-normal, LLR>>1, *p* < 0.005; constant across all recording days). The stability of the power law in size was associated with an average slope α = 1.32 ± 0.05 for all arrays (Table 1), which is close to the expectations for the size exponent of a critical branching process (Fig. 2E, F) (Harris, 1963). These results demonstrate the presence of avalanche dynamics in all NHPs for up to 2 weeks in PM and up to 10 weeks in PF.

**Figure 2:**
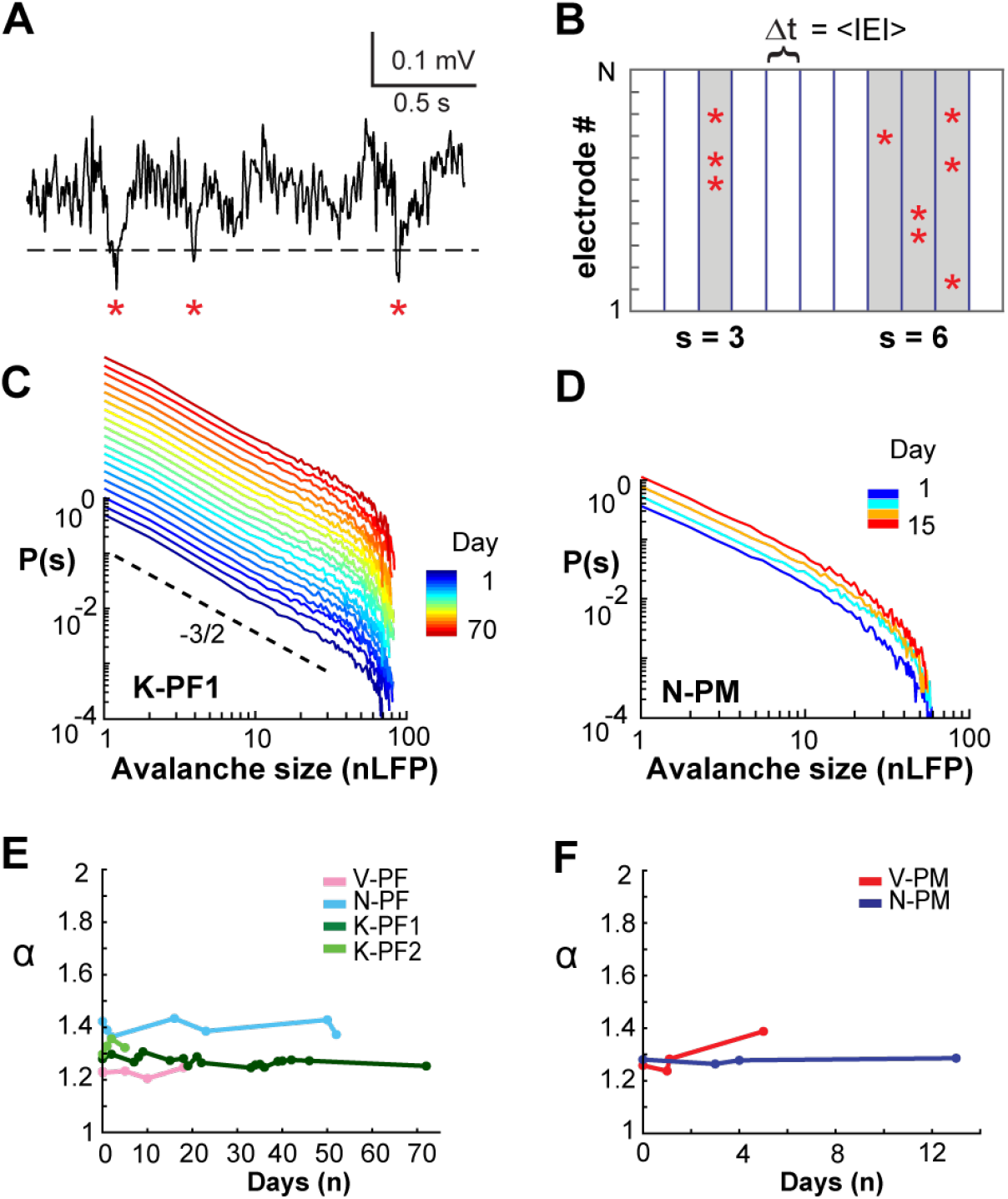
Neuronal avalanche dynamics is stable over weeks in frontal cortex of NHPs. **A,** Sample ongoing LFP at single electrode. A negative threshold (*dashed line*) is applied to identify significant negative deflections from the mean (nLFPs, *red asterisks*). **B,** Schematic nLFP raster at *Δt* = ⟨IEI⟩ showing neuronal avalanches identified as concurrent or consecutive nLFPs on the array separated by at least one empty time bin. Two avalanches (*gray*) of size 3 and 6 are identified. **C, D,** Power law in size *S* for nLFP avalanches for all recordings (*color bar*) in two NHP (*Δt* = ⟨IEI⟩). *Broken line*: power law with slope −3/2 as visual guide. *Color bar*: Recording day. Size distributions remain power laws for all days (LLR values > 0 significant; vs. exponential distribution; see also Table 1). **E, F,** Stability of slope α as a function of time (*color coded*).

### Long-term stability in the temporal profile of avalanches and corresponding scaling collapse

The temporal profile of a neuronal avalanche describes how avalanches unfold in time by initiating locally, growing in magnitude, and eventually contracting spatially before ending. Recent work (Miller et al., 2019) confirmed the prediction that the temporal profile of neuronal avalanches obeys the characteristic motif of an inverted parabola and that empirically-observed mean avalanche profiles collapse onto this scale-invariant motif regardless of avalanche duration. For this analysis, the temporal resolutions were set to *Δt* = 1 ms and *Δt* = 30 ms respectively, which demonstrated stable power law size distributions and expected change in slope α (Table 1; 1 ms: all LLR > 10^4^, all p << 0.005; 30 ms: all LLR > 10, all p << 0.005) as reported previously (Beggs and Plenz, 2003; Petermann et al., 2009). These resolutions allow for the collapse of temporal avalanche profiles in the presence of transient γ–oscillations in ongoing PF and PM activity of NHPs (Miller et al., 2019).

We explored the stability of these scaling predictions over time, finding that the characteristic motif for mean profiles of avalanche lifetime, *L*, with *L* = 3*Δt* – 5*Δt* did trace inverted parabolas for fine (*Δt* = 1 ms, Fig. 3A, B,) and coarse (*Δt* = 30 ms, Fig. 3C, D) temporal resolutions (for statistical comparison see Table 2 for individual arrays). Importantly, the scaling exponent χ producing these parabolic, collapsed motifs was close to the value of 2 expected for a critical branching process (di Santo et al., 2017; Miller et al., 2019; Sethna et al., 2001) and did not change over time (Fig. 3B, D; for *Δt* = 1 ms, χ = 1.93 ± 0.14, linear regression fit of 1*10^−2^ ± 0.2; for *Δt* = 30 ms, χ = 2.01 ± 0.24, linear regression slope of 2*10^−2^ ± 0.04; Table 3 for individual arrays). In contrast, time-shuffled data reliably produced flattened motifs (Fig. 3E, F) with a scaling exponent close to 1, as found for non-critical dynamics, in which sizes grow linearly with duration (Villegas et al., 2019).

**Figure 3:**
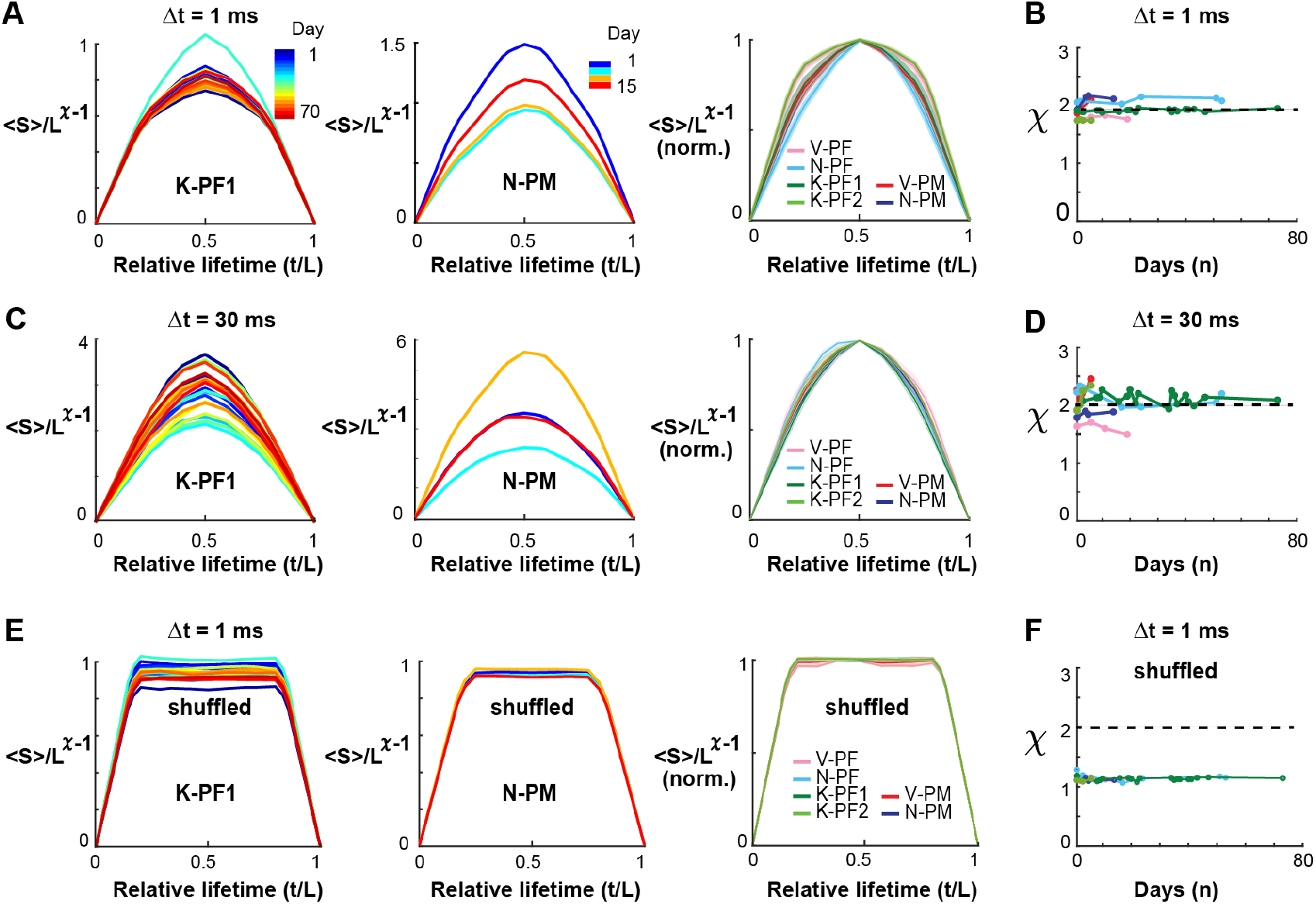
Collapse in the temporal profiles of neuronal avalanches and corresponding scaling exponent χ = 2 are stable over weeks in frontal cortex. **A**, Collapse in the temporal profile of avalanches at *Δt* = 1 ms. Best collapse per recording day (*color bar*) for K-PF1 (*left*) and N-PM (*middle*). Rightmost column summarizes the normalized collapse motif (*norm*.) averaged across days for each NHP and cortical area. Shaded region indicates mean ± standard error per array within line thickness. **B,** The scaling exponent χ at which the best temporal profile collapse is obtained remains close to the theoretical value of 2 predicted for a critical branching process for all recording sessions. **C, D** Same as in *A*, *B* but at *Δt* = 30 ms demonstrating the characteristic parabolic profile for coarse-grained temporal scale. **E**, Same as in *A* – *D*, but nLFP events have been shuffled in time to remove spatial and temporal correlations. Note that the characteristic profile no longer traces a parabola as expected for critical dynamics. **F**, The scaling exponent χ is close to 1 when correlations are destroyed.

**Table 2.** Summary of scaling function estimate for the temporal profile of neuronal avalanches. A parabolic fit (Δ_parab_) is compared to a semicircle fit (Δ_semi_) at temporal resolutions *Δt* = 1 ms and Δt = 30 ms for each nonhuman primate and array and its corresponding phase-randomized profile (*random*) at *Δt* = 1 ms.

| | Temporal Profile Fit:<br>Parabola vs. Semicircle<br>RMSE ($\Delta t = 1$ ms) | | Temporal Profile Fit:<br>Parabola vs. Semicircle<br>RMSE ($\Delta t = 30$ ms) | | Random Profile Fit:<br>Parabola vs. Semicircle<br>RMSE ($\Delta t = 1$ ms) | |
| --- | --- | --- | --- | --- | --- | --- |
|  | Paired t-Test<br>(*p*-value) | ANOVA<br>(F) | Paired t-Test<br>(*p*-value) | ANOVA<br>(F) | Paired t-Test<br>(*p*-value) | ANOVA<br>(F) |
| **V-PF** | 4.36E-05 | 110.44 | 2.00E-04 | 62.26 | 5.96E-10 | 4832.54 |
| **V-PM** | 3.87E-04 | 121.17 | 3.40E-05 | 417.07 | 7.82E-07 | 2776.65 |
| **N-PF** | 6.12E-11 | 460.49 | 2.66E-13 | 1156.15 | 3.74E-14 | 1607.96 |
| **N-PM** | 6.81E-08 | 991.66 | 3.15E-06 | 272.38 | 2.41E-10 | 6533.12 |
| **K-PF1** | 2.76E-22 | 472.2 | 2.41E-25 | 715.09 | 1.98E-50 | 18530.05 |
| **K-PF2** | 2.60E-06 | 290.68 | 9.64E-06 | 186.09 | 1.91E-09 | 3277.87 |
| **Fit Error $\Delta$** | $\Delta_{\text{parab}} < \Delta_{\text{semi}}$ | $\Delta_{\text{parab}} < \Delta_{\text{semi}}$ | $\Delta_{\text{parab}} < \Delta_{\text{semi}}$ | $\Delta_{\text{parab}} < \Delta_{\text{semi}}$ | $\Delta_{\text{parab}} > \Delta_{\text{semi}}$ | $\Delta_{\text{parab}} > \Delta_{\text{semi}}$ |

**Table 3.** Summary of estimates for the scaling exponent χ to optimally collapse temporal avalanche profiles at temporal resolutions Δt = 1 ms and Δt = 30 ms for each nonhuman primate and array. The stability of χ across all recording days was quantified by the linear regression slope reported and 95% confidence limits. *Random*: Phase randomized control of temporal avalanche profile.

| | Scaling<br>Exponent $\chi$<br>($\Delta t = 1$ ms) | $\chi$<br>Stability<br>($\Delta t = 1$ ms) | Scaling<br>Exponent $\chi$<br>($\Delta t = 30$ ms) | $\chi$<br>Stability<br>($\Delta t = 30$ ms) | Scaling<br>Exponent $\chi$<br>Random<br>($\Delta t = 1$ ms) | $\chi$<br>Stability<br>Random<br>($\Delta t = 1$ ms) |
| --- | --- | --- | --- | --- | --- | --- |
| | Mean $\chi$<br>(± SD) | Regression<br>slope<br>(95% Cfl) | Mean $\chi$<br>(± SD ) | Regression<br>slope<br>(95% Cfl) | Mean $\chi$<br>(± SD) | Regression<br>slope<br>(95% Cfl) |
| **V-PF** | 1.78 ± 0.05 | 0.002<br>(-0.019, 0.022) | 1.61 ± 0.09 | -0.01<br>(-0.02, 0.01) | 1.15 ± 0.01 | 0.001<br>(-0.001, 0.003) |
| **V-PM** | 1.93 ± 0.15 | 0.051<br>(-0.3, 0.4) | 2.25 ± 0.20 | 0.06<br>(-0.6, 0.7) | 1.15 ± 0.00 | 0.000<br>(-0.001, 0.002) |
| **N-PF** | 2.10 ± 0.06 | 0.002<br>(-0.000, 0.004) | 2.15 ± 0.15 | -0.00<br>(-0.01, 0.00) | 1.14 ± 0.03 | 0.000<br>(-0.001, 0.002) |
| **N-PM** | 2.05 ± 0.13 | 0.011<br>(-0.053, 0.075) | 1.85 ± 0.05 | 0.005<br>(-0.016, 0.027) | 1.14 ± 0.01 | 0.002<br>(-0.005, 0.008) |
| **K-PF1** | 1.92 ± 0.03 | 0.000<br>(-0.001, 0.001) | 2.11 ± 0.1 | -0.000<br>(-0.003, 0.003) | 1.15 ± 0.01 | 0.000<br>(-0.000, 0.000) |
| **K-PF2** | 1.76 ± 0.02 | 0.007<br>(-0.004, 0.018) | 2.12 ± 0.20 | 0.086<br>(-0.029, 0.2) | 1.15 ± 0.00 | 0.000<br>(-0.004, 0.005) |
| **Mean**<br>**± SD** | **1.93 ± 0.14** | **0.012 ± 0.19** | **2.01 ± 0.24** | **0.02 ± 0.04** | **1.15 ± 0.01** | **0.001 ± 0.001** |

### Persistence of integrative global network properties

In order to quantify in more detail the spatiotemporal correlations underlying persistent avalanche activity, we reconstructed functional networks from neuronal avalanches using the NC reconstruction (Pajevic and Plenz, 2009). This method uses a shortcut to Bayesian estimation of network connections, making it approximate, but also computationally efficient, and hence applicable to reconstructing relatively large networks from lengthy event train recordings (Fig. 4A). It also reduces the influence of indirect common input correlations and has been found in simulations to accurately and efficiently reconstruct the directed networks and weights from simulated point-process dynamics (Pajevic and Plenz, 2009). An example of the directed connectivity matrix for a single recording session derived from spontaneous avalanche activity is shown in Figure 4B. We found that for all arrays and recording sessions, the weights of the directed functional connections were not particularly heavy-tailed and distributed between an exponential and lognormal function over more than 3 orders of magnitude (Fig. 4C – E).

**Figure 4:**
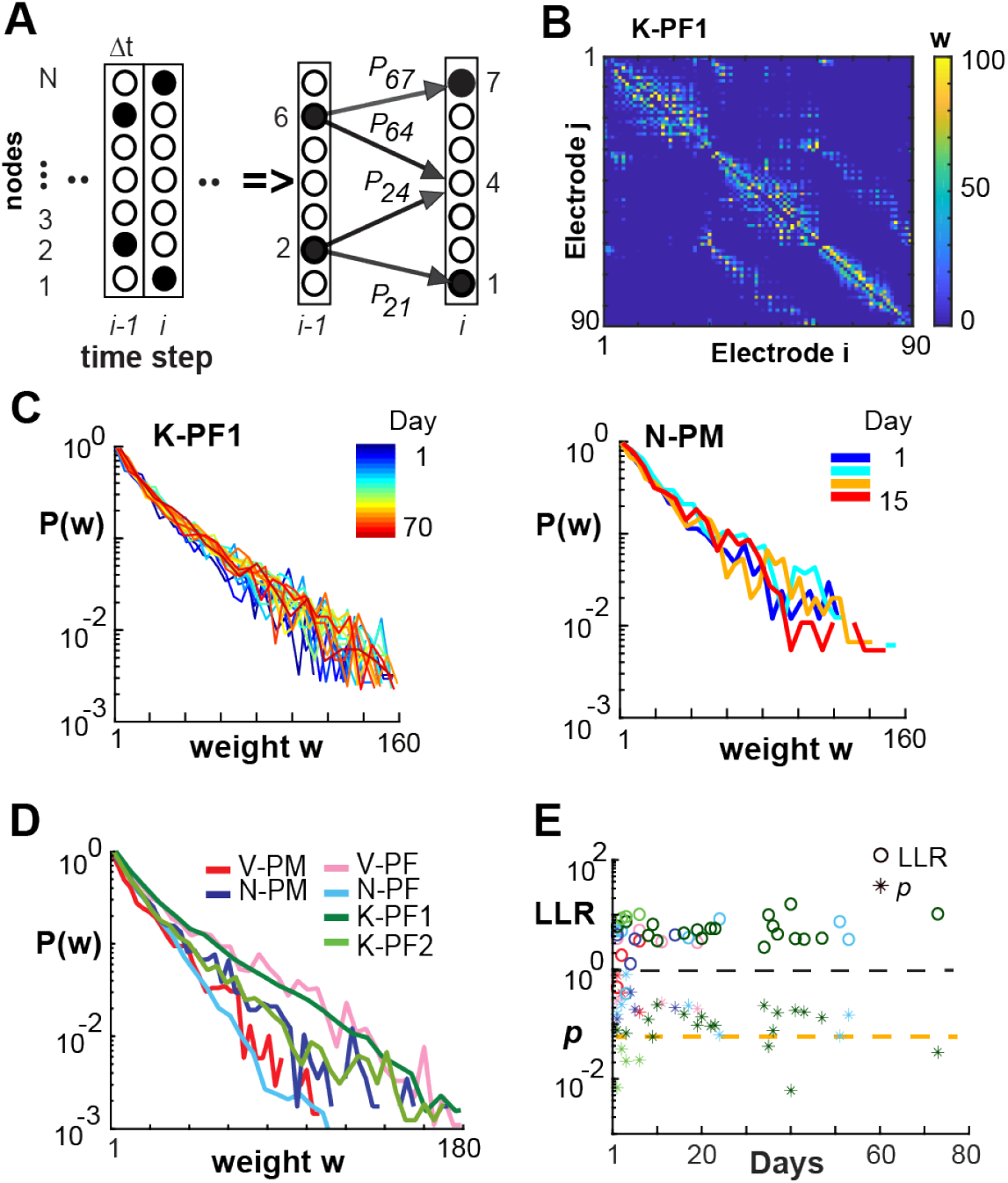
NC reconstruction of directed functional connectivity from ongoing LFPs reveals stable exponential distributions in link weights across time for all NHPs. **A,** Sketch of the NC algorithm. Active nodes (*filled circles*) at time *Δt* _i-1_ are evaluated regarding all existing prior probabilities (*solid* arrows) to obtain the pattern of active and inactive nodes at the next time step *Δt*_i_ (for details see (Pajevic and Plenz, 2009)). **B**, NC reconstructed network based on 15 minutes of nLFP activity at *Δt* = 1 ms from K-PF1 produces an adjacency matrix of weights *w*_ij_ showing sparse and mostly local functional connectivity. **C,** Normalized distributions in directed link strength for all recording days (*left*, K-PF1; *right*, N-PM). *Color bar*: recording day. **D,** Normalized probability distribution trends were consistent across areas and NHPs. Average distribution over all recording days. *Color code*: NHP-area. **E**, LLR (*open circles*) and *p* values (*asterisks*) for exponential vs. log-normal comparison of daily weight distributions (e.g. *C*) for all arrays and days (color code as in *D*).

Having reconstructed the directed, weighted links in our avalanche-derived functional networks, we proceeded to calculate several graph theoretical properties. Integrative networks are characterized by a positive correlation, *R*_*CL*_, between the link cluster coefficient, *C*_*L*_, and the corresponding link weight, *w*, between two nodes (Fig. 5A; (Pajevic and Plenz, 2012)). This positive correlation demonstrates that strongly connected nodes have more common neighbors compared to weakly connected nodes (Fig. 5B). We employed degree-sequence preserved randomization to subtract potential contributions to link clustering from the node degree distribution and report here the resulting excess link clustering, *ΔC*_*L*_ (see Material and Methods). The monotonic increase in *ΔC*_*L*_ with lowest-to-highest weight rank (Fig. 5C) clearly identifies integrative functional network organization in both PM and PF that was stable over time (Fig. 5D; linear regression slope of 5 ± 10 ×10^−3^; 95% confidence limit; see Table 4 for individual arrays).

**Figure 5:**
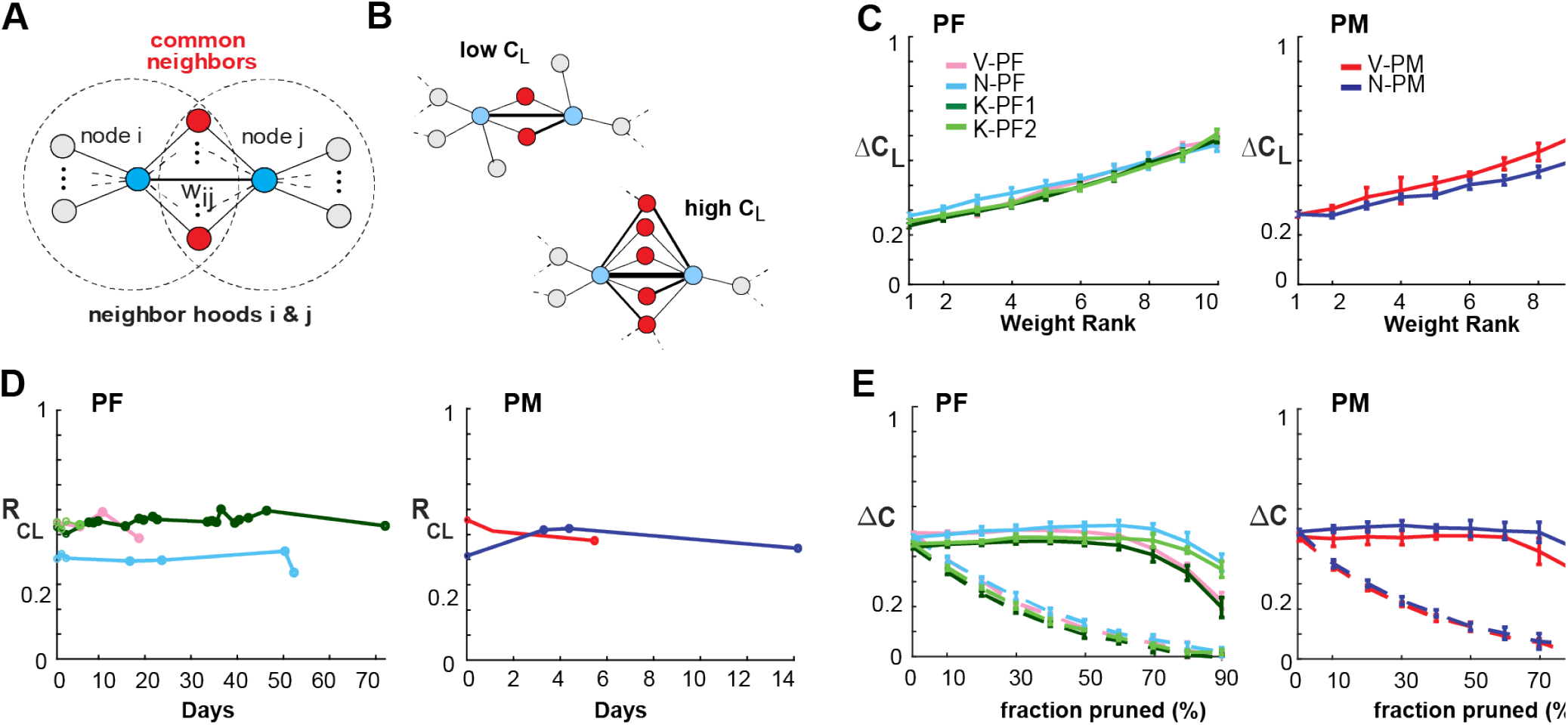
Link-clustering analysis reveals that integrative global network properties are stable over time. **A,** Schematic clustering diagram depicting two nodes (*blue*) connected by a link with weight *w*_*ij*_. In an integrative network, the proportion of common neighbors (*red*) is expected to positively correlate with *w*_*ij*_. **B,** Sketch of two subnetworks with low and high link clustering *C*_*L*_ respectively. **C,** The excess link clustering Δ*C*_*L*_ (see Methods) increases with link strength in all prefrontal (*left*) and premotor (*right*) cortical arrays (*color coded*; NHP-area). **D,** Analysis of cortical networks showed strongly positive *R*_*CL*_ over all recording sessions for up to 73 days. **E,** Link-pruning analysis of excess node clustering *ΔC* calculated via degree-sequence preserved randomization (Pajevic and Plenz, 2012) showed that the integrative clustering motif was robust to weak-link pruning (*solid lines*; error bars denote mean ± SD over all recordings for an array), while the clustering “backbone” of the network quickly degraded when strong links were pruned first (*dotted lines*; *color code* as in *D*). *C* – *E*, *Δt* = 1 ms.

**Table 4.**
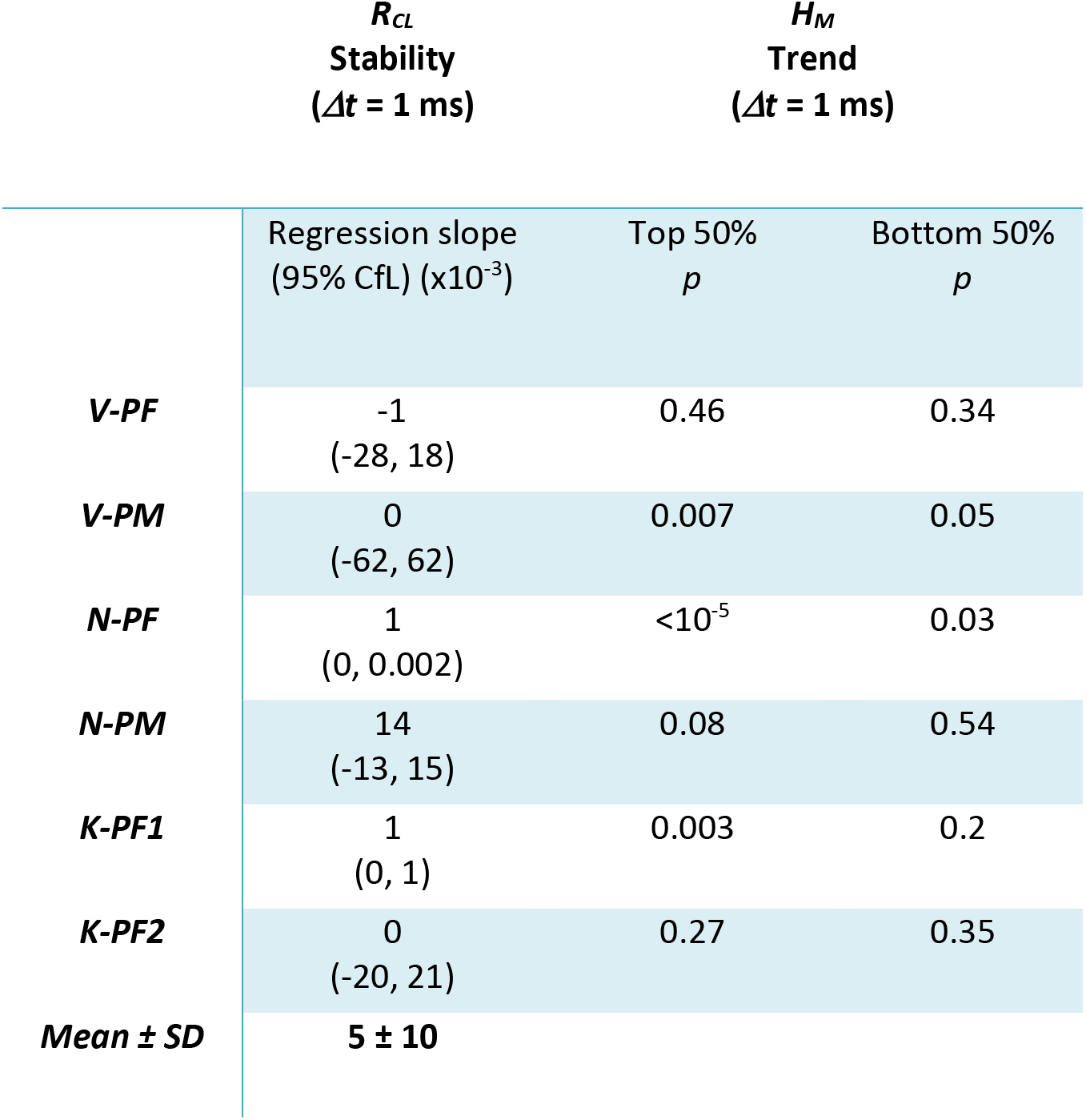
Functional network stability based on NC reconstruction algorithm at Δt = 1. Shown is the stability of *R*_*CL*_ over time (see also Fig. 5D). H_M_ *p*-values given for top 50% and bottom 50% for significant trend in mixing observed.

Pruning analysis is an efficient approach to study the weight organization of networks (Onnela et al., 2007; Pajevic and Plenz, 2009). We found that pruning from the bottom, i.e. pruning weakest links progressively, had negligible effect on the network clustering properties. This was evidenced by a near constant value of the excess node clustering, *ΔC,* until the vast majority (80 – 90%, Fig. 5E; solid lines) of links were removed. In contrast, pruning from the top, i.e. removing the strongest links first, rapidly decreased *ΔC*, a result consistently found for all NHPs (Fig. 5E). We conclude that the integrative property arises from a “backbone” of strong links that is robust to weak-link pruning, yet quickly degrades when strong links are removed.

### Entropy of mixing identifies link progression over many weeks

Given the stability observed over weeks in the global network dynamics, i.e., <*R*>, α, χ, and integrative network organization, i.e., *R*_*CL*_, we next sought to analyze in more detail the stability at the local level of individual link weights. Specifically, we examined whether the stability identified at the global level simply arises from stability at the local level. To quantify changes at the local level, we categorize the links, e.g., weak, medium, and strong, and analyze whether links switch between categories over time, e.g., from weak to medium or vice versa. As a measure of mixing, we calculate the entropy of mixing, *H*_*M*_, (see Methods). Two main factors contribute to potential changes in *H*_*M*_, which originate from changes in link categories in a network from one day to another. First, independent random changes on each day will result in increased mixing and *H*_*M*_, but not progressively over time. For example, errors in network reconstruction alone will result in a constant *H*_*M*_ over many days if the underlying network is not changing. Second, link weights could gradually evolve over days and weeks, and potentially change their categories, which is now progressively mixed and therefore is expected to lead to a monotonic increase in *H*_*M*_ until the maximum in mixing is reached. Because reconstruction errors are higher for weak links compared to strong links, we quantified potential changes in link weight from NC reconstructed networks separately for the strongest (top 50%) and weakest (bottom 50%) links. This separation is also motivated by our results for top- vs. bottom-pruning, which demonstrated that networks are less robust to link changes in the top half of the link weight distribution. For each half, we partitioned link weights further into strength quintiles, yielding 5 categories, and then studied whether members from different categories will mix over time. In figure 6, we show the data for the top half and bottom half and compare those curves to *H*_*M*_ obtained if the category labels were fully randomized at each step and also present the theoretical maximum for *H*_*M*_ (see also Material and Methods). Because all successive distributions used the link labels derived from Day 1 (making *H*_*M*_ = 0), the average entropy of mixing was plotted beginning from the second recording for each NHP. For all arrays, we found that *H*_*M*_ for the bottom half of links reaches close to the maximal entropy by the following day demonstrating that weak links fluctuate randomly. On the other hand, we found progressive and statistically significant increases of *H*_*M*_ over subsequent days for strong links in three out of six arrays (Figure 6A, B; see Table 4). These findings demonstrate stability in avalanche dynamics and integrative network organization even though the network undergoes significant weight reorganization at the individual link level.

**Figure 6:**
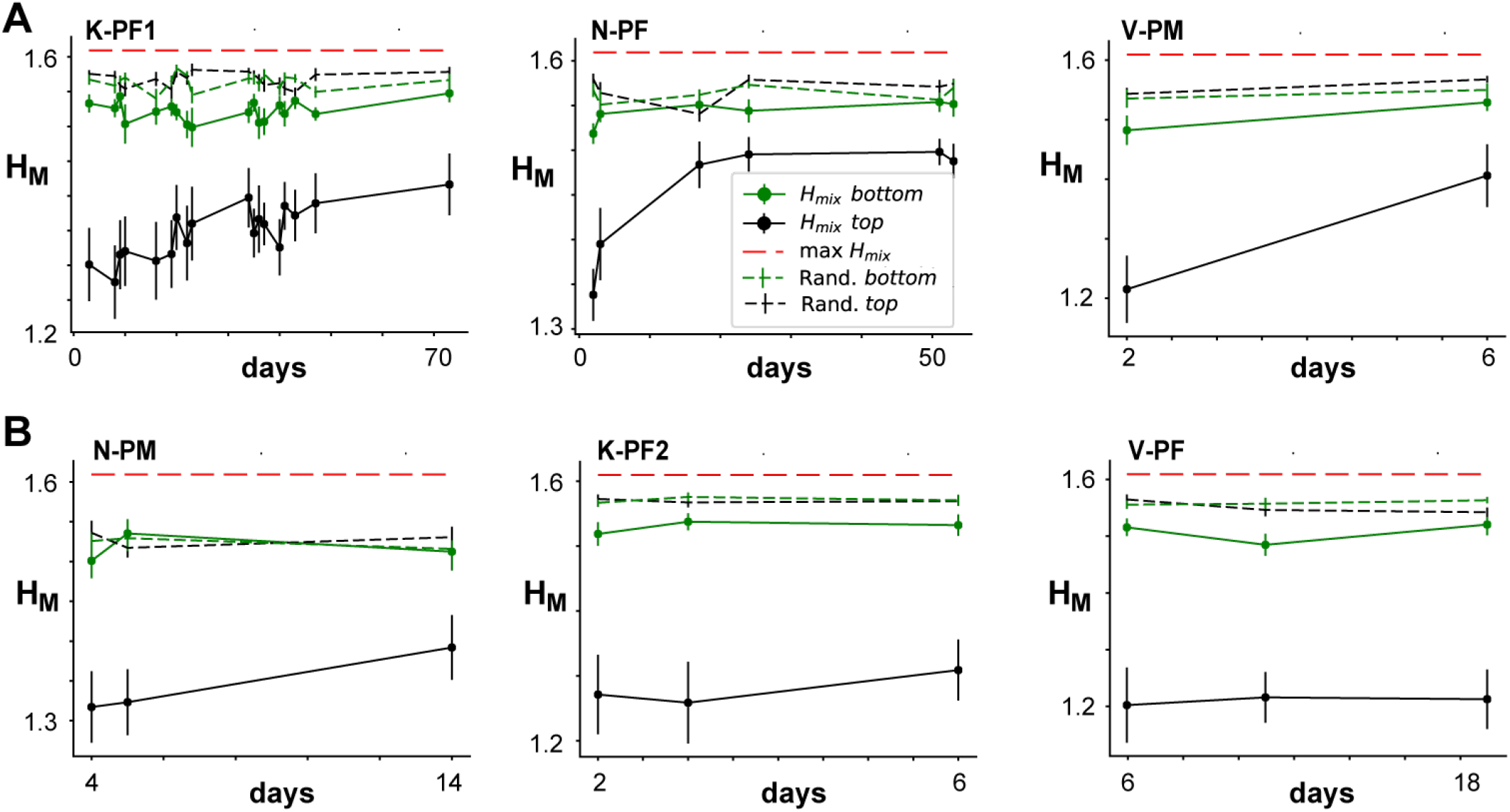
Entropy of mixing analysis demonstrates progressive changes in strong links over time for some networks. Entropy of mixing (*H*_*M*_) analysis of the link weights obtained with NC reconstructions from Δt = 1 ms raster recordings. *H*_*M*_ is plotted for the “top” half (strongest 50% of links; *black curves*) and “bottom” half (weakest 50% of links; *green curves*) versus the days after the initial recording. The black and green dashed lines indicate corresponding *H*_*M*_ obtained when the labels were fully randomized, while the red dashed line indicates the theoretical maximum of *H*_*M*_. The six panels represent six different arrays from the three monkeys shown in two rows. **A**, Arrays for which statistically significant progression of mixing was obtained for “top” links. **B**, Arrays for which no statistically significant progression was observed. The “top” half of the links consistently showed lower *H*_*M*_ than the “bottom” half of the links, which is mainly due to the smaller reconstruction error. The Day 1 points are not plotted because, by default of our labeling, the initial value for *H*_*M*_ is zero.

## Discussion

Fluctuations in ongoing neuronal activity give rise to functional networks that allow for the categorization of healthy and pathological brain states (Bullmore and Sporns, 2009; Fox and Raichle, 2007; Raichle, 2015), yet, the dynamical and graph theoretical markers underlying these ongoing fluctuations have been challenging to identify. By recording the LFP in prefrontal and premotor cortex of NHPs over weeks at high spatiotemporal resolution, we established two robust markers of ongoing activity; the dynamical category of neuronal avalanches and the graph theoretical category of integrative networks. Avalanche dynamics revealed power law statistics, parabolic temporal profiles and a scaling exponent of 2 in support of critical branching process dynamics (di Santo et al., 2017; Miller et al., 2019; Sethna et al., 2001). We further demonstrate that avalanches form integrative networks despite reorganization of individual link weights as identified using NC estimators and an entropy of mixing approach (Onnela et al., 2007; Pajevic and Plenz, 2009, 2012). Our results suggest that avalanche dynamics and corresponding integrative networks identify a robust state of the awake frontal cortex. This work should be impactful for computational studies of cortex function and for translational research into the identification of brain dynamics in healthy subjects and corresponding deviation in brain disorders.

Recent analysis of evoked cortical activity in nonhuman primates during behavior demonstrated the preservation of neuronal avalanche dynamics in the face of transient activity changes, specifically the nLFP rate (Yu et al., 2017). Given the robustness of integrative network properties observed in combination with avalanche dynamics *in vitro* (Pajevic and Plenz, 2009, 2012) as well as *in vivo* (the present study), we would expect integrative networks properties to also persist during behavioral tasks.

### Stability of LFP based avalanches and relation to neuronal activity

Technical advances in identifying long-term robustness in neuronal activity using cellular resolution approaches (Dickey et al., 2009; Fraser and Schwartz, 2011; Jackson and Fetz, 2007; Li et al., 2017; Nicolelis et al., 2003; Tolias et al., 2007) typically suffer from spatial and temporal subsampling (Levina and Priesemann, 2017; Petermann et al., 2009; Ribeiro et al., 2010; Ribeiro et al., 2014) limiting proper identification of avalanche dynamics and corresponding reconstruction of functional connectivity. The LFP, on the other hand, is a robust neuronal population signal, less prone to spatial subsampling and at high temporal resolution contains local spike information particularly when recorded from superficial layers (Donoghue et al., 1998; Mehring et al., 2003; Petermann et al., 2009; Rasch et al., 2008). Indeed, we recently demonstrated in nonhuman primates and *in vitro* slices that single neurons selectively participate in expansive, repeated LFP avalanches (Bellay et al., 2021), which suggest that the stability observed in the present study for LFP-based avalanches might extend to the single neuron level.

Experimental manipulations support a homeostatic regulation of avalanche dynamics in cortex. Transient pharmacological perturbation *in vitro* (Plenz, 2012) or sensory deprivation *in vivo* (Ma et al., 2019) initially abolishes avalanche dynamics, which is followed by a recovery over several days. Developmental findings further support an autonomous regulation of neuronal avalanches in isolated cortex *in vitro* in the absence of sensory input (Pasquale et al., 2008; Stewart and Plenz, 2007; Tetzlaff et al., 2010). Identifying the homeostatic regulation of avalanche dynamics and integrative networks will require precise perturbation approaches (Chialvo et al., 2020) in combination with advanced cellular techniques suitable for long-term monitoring such as ratiometric genetically-encoded calcium indicators (GECIs), which naturally compensate for cell-intrinsic expression variability (Lutcke et al., 2013).

### NC reconstruction of integrative networks

Reconstruction of our directed, weighted functional networks utilized a Bayesian-derived NC approach, which is easily scalable to large networks (Pajevic and Plenz, 2009, 2012) and reduces correlations from common input. Network simulations demonstrated ground-truth reconstruction for various small-world topologies from subcritical, supercritical and critical dynamics, the latter in line with reconstructions from observed avalanche activity (Pajevic and Plenz, 2009).

Here we used significant events in LFP fluctuations, i.e. nLFPs, for the reconstruction of functional connectivity, which differs from the more common ansatz based on continuous timeseries. Accordingly, our approach, which selectively reconstructs synchronization dynamics in the form of avalanches, introduces a time scale Δt to which the NC reconstruction is sensitive. In order to identify the spatiotemporal spread of avalanches, the use of a microelectrode array introduces a spatial discretization, which consequently enforces a discretization Δt in time. It was found empirically that Δt should be linked to the average propagation velocity of neuronal activity in the network, allowing Δt to be approximated by <IEI> (Beggs and Plenz, 2003; Petermann et al., 2009). For Δt smaller than <IEI>, errors of prematurely terminating avalanches increase, whereas errors of concatenating successive avalanches increase for Δt larger than <IEI>. An approximate balance regarding these two errors is struck for Δt = IEI. With respect to NC reconstruction, increasing Δt will increase the count in nLFP occurrences on the array per time step, resulting in an increase in the computational load and increased error in estimating priors for potentially activated links. Accordingly, we chose a Δt slightly smaller than IEI for a conservative and balanced estimate of the functional connectivity based on avalanche propagation.

Integrative networks differ from networks in which the weight is positively correlated with the node degree (Barrat et al., 2004; Bianconi, 2005; Pajevic and Plenz, 2012). Our shuffling correction further demonstrates that the organization of weighted links in integrative networks cannot be simply explained by the link-degree distribution and in which strong links are part of a continuous, in our case near exponential distribution (*cf*. Fig. 4D). Integrated networks are also not a simple consequence of critical branching processes generating avalanches, which can be realized in many topologies and architectures (Pajevic and Plenz, 2009, 2012).

### Robust avalanche scaling of χ = 2 over many weeks

Heterogeneous activity in the form of neuronal avalanches has been the hallmark of ongoing (Beggs and Plenz, 2003; Bellay et al., 2015) and evoked (Yu et al., 2017) brain activity in superficial layers of the cerebral cortex. The scale-invariant hallmark of avalanches has been found to capture ongoing neuronal activity in nonhuman primates at the mesoscopic level in the LFP (Klaus et al., 2011; Petermann et al., 2009; Yu et al., 2014; Yu et al., 2017; Yu et al., 2011) and macroscopic activity measured with EEG (Meisel et al., 2013), MEG (Shriki et al., 2013) and fMRI (Fraiman and Chialvo, 2012; Tagliazucchi et al., 2012). The observed power law distributions in cluster size and duration support the idea that the cortex operates close to a critical state at which networks gain numerous advantages in information processing (Beggs and Plenz, 2003; Gautam et al., 2015; Kinouchi and Copelli, 2006; Shew et al., 2009; Shew et al., 2011; Yang et al., 2012).

Our present demonstration of an inverted-parabolic avalanche shape, collapse exponent of 2 and size distribution slope >−2 is in line with predictions that the unfolding of avalanches in the adult brain is governed by a critical branching process (for details see (Miller et al., 2019)). This profile demonstrates that the initial spatial unfolding of an avalanche in time is truly expansive and should not be viewed as being dominated by a few strong links. Recently, avalanches with a scaling factor of 2 have been described for whole-brain neuronal activity in the zebrafish larvae (Ponce-Alvarez et al., 2018), however, the corresponding size and duration exponent were significantly steeper compared to reports for the mammalian cortex, suggesting a different dynamical model for the zebrafish.

Connectome analyses on a time scale of seconds have identified transient fluctuations in functional connectivity reflecting distinct states (Hutchison et al., 2013; Liu and Duyn, 2013; Zalesky et al., 2014), which based on computational modeling were found to be intermittent, a signature of metastability (Deco et al., 2017; Hansen et al., 2015). Because our analysis was based on nLFPs and not continuous time signals, we were unable to obtain NC reconstructed networks for shorter time periods than 15 – 30 min. In fact, our entropy of mixing analysis allowed for an assessment of link weight progression in the face of reconstruction errors due to limited data availability. Transient fluctuations in functional connectivity estimates are not in contrast to the reorganization of links reported in our study. Short-term fluctuations in the face of long-term stability are hallmarks of critical dynamics (Fraiman and Chialvo, 2012; Tagliazucchi et al., 2012).

### Potential advantages of avalanche dynamics and integrative networks for cortex function

Anatomically, the mammalian cortex is a sheet of local, functional modules that dynamically combine to support complex brain functions (Braitenberg and Schüz, 1991; Honey et al., 2007). Local connectivity ensures diverse local operations whereas long-range connections support global coordination (e.g. (Bullmore and Sporns, 2009; Kirst et al., 2016)). These structural hallmarks are dynamically realized in neuronal avalanches, which support a scale-free and selective organization of neuronal synchronization over all distances. They are graph theoretically embedded in integrative networks, which maintain a high level of node clustering until about 90% of the strongest links are pruned, i.e. about 10% of strong links are used for long-range connections. Intriguingly, simulations identified a particular local learning rule that operates on activity cascades such as avalanches to build integrative networks from random networks (Pajevic and Plenz, 2012). The rule increases link weights at locations of recent cascade failure, thereby facilitating the unfolding of future avalanches (Alstott et al., 2015). This mechanism opens bottlenecks in the network and further supports activity propagation over long distances. We therefore hypothesize that avalanche dynamics in combination with integrative network organization are beneficial for local operational diversity while supporting long-range, selective propagation of information.

## Material and Methods

### Animal procedures

All animal procedures were conducted in accordance with NIH guidelines and were approved by the Animal Care and User Committee of the National Institute of Mental Health. Three adult NHPs (*Macaca mulatta*; 1 male, 2 females; 7–8 years old) received two chronic implantations of high-density 96-microelectrode arrays each (Blackrock Microsystems; 4×4 mm^2^; 400 μm interelectrode distance; 10×10 grid with corner grounds). To direct recordings towards superficial cortical layers II/III, electrode shanks of 0.6 mm length were used in prefrontal cortex (PF; n = 4), and shanks of 1 mm length were used for premotor cortex (PM; n = 2). During recording sessions monkeys sat head-fixed and alert in a monkey chair with no behavioral task given. Portions of this dataset have been analyzed previously (Bellay et al., 2021; Meisel et al., 2013; Miller et al., 2019; Yang et al., 2012; Yu et al., 2014).

### Local field potential recordings in awake monkeys

Simultaneous and continuous extracellular recordings were obtained for 12 – 60 min per recording session (2 kHz sampling frequency), band-pass filtered between 1 – 100 Hz (6th-order Butterworth filter) to obtain the local field potential (LFP), and notch-filtered (60 Hz) to remove line noise. About 2 ± 1% of time periods were removed from functional electrodes due to artifacts introduced by e.g. vocalization, chewing, sudden movements. These artifacts were identified by threshold crossing (SD > 7) and excised (± 0.25 s). Arrays on average contained 86 ± 8 functional electrodes that exhibited 64 ± 50 μV of spontaneous LFP fluctuations (SD). Channels which had been removed from an array at any recording day were discounted for all recording days. Electrode LFPs were z-transformed and recording sessions for each array were analyzed individually. The current study represents a combined 26 hr of ongoing cortical LFP activity.

### Neuronal avalanche definition and temporal resolution

For each electrode in the array, peak amplitude and time of negative LFP (nLFP) threshold crossings (−2 SD) were extracted at the temporal resolution of *Δt* = 0.5 ms given our sampling rate of 2 kHz. We note that the negative peak amplitude of the LFP correlates with the probability of extracellular unit firing and synchrony in nonhuman primates (Bellay et al., 2021; Petermann et al., 2009). The mean inter-event interval, <IEI>, defined as the average time between consecutive nLFPs among all functional electrodes on the array, was calculated. We then binned nLFP times in steps of *Δt* = <IEI> for each electrode, which ranged between 3 – 4 ms across NHPs and arrays. All nLFP events from all electrodes were combined into a matrix, i.e. raster, with rows representing electrodes and columns representing time steps. A population time vector was obtained by summing nLFPs in the raster for each time step. Avalanches were defined as spatiotemporal continuous activity in the population vector bracketed on each side by at least one time bin of duration *Δt* with no nLFP. The size of an avalanche, *S*, was defined as the number of nLFPs participating. Multiple nLFPs at an electrode during an avalanche are rare (Yu et al., 2014) and were counted in size estimates. Scale-invariance of *S* was visualized by plotting probability distributions P(S) in double-log coordinates. We previously showed that analyzing nLFP at *Δt* = <IEI> results in a distribution of avalanches with a slope close to −3/2 in line with expectations for a critical branching process (e.g., (Beggs and Plenz, 2003)).

The maximum log-likelihood ratio (LLR) was calculated to test potential power law distributions in avalanche size *S* against the alternatives of exponential or log-normal distribution models (Clauset et al., 2009; Klaus et al., 2011). When tested positive for power law, the LLR estimate for best slope α was reported. Here we used the range of *S* = 1 to 40, which excludes the distribution cut-off (Klaus et al., 2011) close to the total number of functional electrodes on the array (n >70). We note that analyzing avalanche dynamics at different temporal resolutions, e.g. at shorter *Δt* or longer *Δt* compared to the <IEI>, increases or decreases the slope α respectively, but not the power law form itself (see also Table 1; (Beggs and Plenz, 2003, Petermann, 2009 #4926)).

### Avalanche temporal profile collapse and scaling exponent

Given the difficulty of obtaining robust lifetime distributions in the presence of ongoing oscillations, the stability analysis for avalanches in the present work focused mainly on the calculation of size distributions and temporal profile collapse (for details see Miller, 2019 #45}. The temporal profile, *S*(*t*), was described by the number of participating electrodes at each timestep t = 1 up to the avalanche lifetime, *L*, which was defined by the number of timesteps *Δt* that the avalanche persisted. In line with our previous results, profile analysis was carried out at *Δt* = 1 ms and *Δt* = 30 ms to avoid profile modulation by ongoing γ-oscillations (Miller et al., 2019). Avalanches were grouped by *L* in multiples of *Δt* and averaged to obtain the mean temporal profile for a given lifetime, ⟨*S*⟩(*t*, *L*). After normalizing to dimensionless time units, i.e., the relative lifetime, *t/L*, amplitudes were then rescaled via Equation 1.

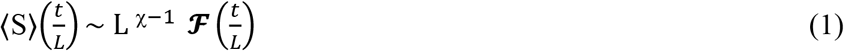

The profile collapse function, shown in Equation 1, relates the mean profile for each lifetime *L*, ⟨*S*⟩(*t/L*), with a characteristic temporal motif 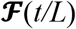, and scaling factor, 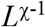, which is independent of *L* according to Equation 1. To perform a shape collapse, we plotted ⟨*S*⟩(*t*, *L*) from *L*−1 through *L*+1 for *L*_min_ = 4*Δt* (to reduce finite size effects in shape caused by too few data points). The collapse error, *Δ*_*F*_, was quantified via a normalized mean-squared error (NMSE) of height-normalized individual profiles to the combined normalized average of all collapsed profiles, 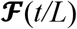. Minimized collapse error was calculated by scanning through χ = 0.5 to 3 at resolution of 0.001 to find the collapse in avalanche waveform associated with the smallest *Δ*_*F*_ via χ_collapse_. A value of *Δ*_*F*_ > 1 was considered a failure in collapse.

We note that for fractal objects such as spatiotemporal avalanches, changing the temporal resolution also partitions avalanches differently. For a *Δt* = 1 ms, avalanche durations in multiples of *Δt* will exhibit a power law as well as for *Δt* = 30 ms (see e.g. (Miller et al., 2019)). Accordingly, the temporal profile of avalanches must be examined in multiples of generations of duration *Δt*. A comparison between timescales in absolute times may not be possible. In the present study, an avalanche of duration 150 ms encompasses L = 5 generations at *Δt* = 30 but would require L = 150 generations at *Δt* = 1, which is highly unlikely to occur.

### Fit of the average temporal profile of avalanches

For the parabolic fit we used the approach by (Laurson et al., 2013) as follows:

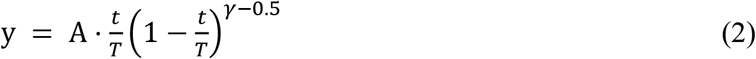

The parabolic fit error, Δ_parab_, was quantified via a normalized mean squared error (NMSE) of individual profiles to an amplitude-matched parabola which was coarse-grained to match *L*. Comparison to a semicircle fit was conducted in the same manner to obtain Δ_semi_ using

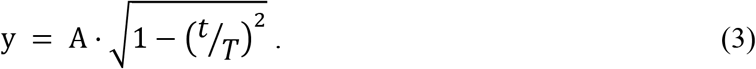

### Correlation-based network reconstruction

Networks were reconstructed for each recording session individually and the links with low pairwise correlations were removed before subsequent network analysis. Rather than applying a strict minimum correlation threshold for removing the links, which would leave our analysis vulnerable to inter-NHP differences in electrode impedances, etc., we instead sequentially removed the weakest links in 0.01 increments of the correlation threshold until a predefined sparsity of approximately 40% was achieved. Weight bins were divided into deciles for pruning analysis (Fig. 4).

### Normalized Count reconstruction of network

To remove the influence of common input to electrodes on the array, which inflates the Pearson’s pairwise correlation, we conducted a network reconstruction. This nonparametric method, called the NC approach, is used for the network reconstruction because its computational efficiency is comparable to the correlation analysis yet with much lower reconstruction error, which are important considerations for analysis of long LFP time series collected in awake *in vivo* preparations. For this reconstruction, the time-window within which active nodes can influence each other was examined for 0.5, 1, <IEI>, and 30 ms. For each node activation (spike), all nodes that were active within Δt, prior to the spike, were deemed as potential causes. Results were very similar for *Δt* = 0.5 and 1 ms (data not shown). For longer *Δt*, the precision of the algorithm decreases rapidly due to (1) the increase in ambiguous potential associations between the number of active nodes in successive frames and (2) a decrease in the number of transition time steps to estimate link weights (Pajevic and Plenz, 2009). Here we show results for *Δt* = 1 ms, which matches our avalanche scaling analysis.

### Integrative networks and Pruning analysis

Once the networks were defined, we studied their integrative properties (Pajevic and Plenz, 2012). The link clustering coefficient, *C*_*L*_, was defined for each link as the ratio of the number of common neighbors of its end nodes to the number of all neighbors (see Fig. 4A). In an integrative network, there is a positive correlation *R*_*CL*_ between *C*_*L*_ and the weight of the link, indicating that strongly connected nodes are likely to share more common neighbors. To visualize this correlation, we divide links based on their weight into equally sized 10 blocks ranging from the weakest block (rank 1) to the highest rank. For each block, we obtain the average *C*_*L*_ and subtract the equivalent value that is obtained from the degree preserved randomized network, yielding the excess link clustering, Δ*C*_*L*_. We also conducted a “pruning analysis” in which we studied the robustness of the excess node clustering ΔC upon the progressive removal of the weakest (bottom pruning) or strongest (top pruning) links. Similarly, the excess node clustering, Δ*C*, was defined as the difference of the mean clustering coefficient of the original network and the corresponding randomized network after the degree-sequence preserved randomization.

### Entropy of mixing analysis

The estimated link weights fluctuated over time in both pairwise correlation and in directed networks obtained with the NC reconstruction algorithm. We sought to explore whether these fluctuations predominantly represent an error in the network reconstruction or arise from a genuine and progressive change in the underlying network weights. To answer this question, we developed a novel method that utilizes the entropy of mixing to quantify the progression of these fluctuations in individual link strengths from one day to another. The entropy of mixing is a concept from thermodynamics and describes the increase in total entropy when partitioned (pure and equilibrated) sub-systems are allowed to mix. It is quantified using the Shannon entropy in which probabilities are replaced by the fractions of each of the original species found at different partitions. Hence, to conduct our entropy of mixing analysis, we need to partition the links into different categories. Labels are assigned according to the magnitude of their weights obtained from the Day 1 data, and partitioning them into a finite set of categories, *N*_*c*_, ranging from the weakest links to the strongest links. In our case, we used *N*_*c*_ = 5, so that the label *c = 1* marked the weakest 20% of the sorted links and incrementally, c = 5 marked the strongest 20% of the links.

We conducted the mixing analysis on two different and complementary sub-networks. The first is the “top” network, consisting of the 50% of the strongest links, and the second is the “bottom” network, consisting of the weakest 50% percent of links. This was done mainly to have some test of the performance of the entropy of mixing analysis, as we expect that the “bottom” half is more dominated by the reconstruction errors of the NC algorithm than the “top” half. Note that with this two-stage stratification of the links, the strongest 20% links in the “top” network are effectively representing the 10% of strongest links in the full network, while the weakest 20% of the “top” network links have still greater magnitude than the strongest quintile of the “bottom” networks, and, *vice versa*, the weakest 20% of the links in the “bottom” represent the 10% of weakest links in the full network. Once the chosen links were labelled, we then monitored the changes of their weights over subsequent days. Specifically, we sort them again based on their new weights, but now partition them over a potentially larger number of weight groups, *Ng = k N*_*c*_, where *k* is an integer, in order to make all groups contain a single category label for the Day 1 stratification (we use k = 3 and *N*_*g*_ = 15, meaning that the links in the groups 1 through 3 will all initially carry the label c = 1, the groups g = 4 − 6 will be labeled c = 2, etc). Due to (1) reconstruction errors and (2) potential actual network weight changes from one day to another, each of the groups will on the subsequent days contain a mixture of the original labels, although it is expected that the low group indices, *g*, will early on predominantly carry the labels of the weak links, and *vice versa*. For each group, we calculate a fraction, *f*_*c*_, of the links that carry label *c*, which by definition are normalized to 1, i.e., 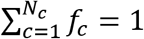. For each group *g* we then calculate the Shannon entropy of such mixture according to 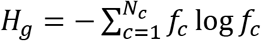. Finally, we report the average and the standard error over all non-edge *Hg* values, i.e., excluding the extreme groups, g = 1 and g = 15, as those have only uni-directional mixing 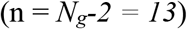. Hence, the average entropy of mixing reported in our results is 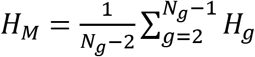.

To evaluate if we have seen a progression in the weight evolution, we test it against the null hypothesis that there was no significant linear trend across the subsequent days. To do so we use a Monte Carlo (MC) procedure, in which we generate one million normally-distributed replicates, N(0,σ_G_) for each point, where σG are the standard deviations of the means obtained using the non-edge groups described in the previous paragraph. For each of the *N*_*MC*_ = 1000000 replicates we perform a linear regression and count the number of times the MC slopes were greater than the slope we obtained using the mean values for *H*_*M*_. From this count, *N*_pos_, our estimate of the *p* values is *p* = (*N*_pos_+1)/(*N*_*MC*_+1) (see Table 4).

## Acknowledgements

We thank members of the Plenz lab and attendees of the “*Critical brain dynamics: 5*^*th*^ *International Workshop on Criticality and the Brain*”, 2016, NIH, USA for their support and useful discussions.

## Author Contributions

S.Y. collected the data; S.R.M. , S.Y. and S.P. analyzed the data; S.R.M., S.Y., S.P. and D.P. wrote the manuscript.

## Supplementary Information

No Supplementary Information.

## References

Alstott, J., Pajevic, S., Bullmore, E., and Plenz, D. (2015). Opening bottlenecks on weighted networks by local adaptation to cascade failures. Journal of Complex Networks 3, 552–565.

Barrat, A., Barthelemy, M., Pastor-Satorras, R., and Vespignani, A. (2004). The architecture of complex weighted networks. Proc Natl Acad Sci U S A 101, 3747–3752.

Bassett, D.S., and Sporns, O. (2017). Network neuroscience. Nat Neurosci 20, 353–364.

Beggs, J.M., and Plenz, D. (2003). Neuronal avalanches in neocortical circuits. J Neurosci 23, 11167–11177.

Bellay, T., Klaus, A., Seshadri, S., and Plenz, D. (2015). Irregular spiking of pyramidal neurons organizes as scale-invariant neuronal avalanches in the awake state eLife 4, e07224.

Bellay, T., Shew, W.L., Yu, S., Falco-Walter, J.J., and Plenz, D. (2021). Selective participation of single cortical neurons in neuronal avalanches. Frontiers in Neural Circuits 14.

Bianconi, G. (2005). Emergence of weight-topology correlations in complex scale-free networks. EL 71, 1029–1035.

Braitenberg, V., and Schüz, A. (1991). Anatomy of the Cortex: Statistics and Geometry (Berlin Heidelberg New York: Springer-Verlag).

Bullmore, E., and Sporns, O. (2009). Complex brain networks: graph theoretical analysis of structural and functional systems. Nat Rev Neurosci 10, 186–198.

Chialvo, D.R., Cannas, S.A., Grigera, T.S., Martin, D.A., and Plenz, D. (2020). Controlling a complex system near its critical point via temporal correlations. Scientific Reports 10, 12145.

Clauset, A., Shalizi, C.R., and Newman, M.E.J. (2009). Power-law distributions in empirical data. SIAM Rev 51, 661–703.

Deco, G., Kringelbach, M.L., Jirsa, V.K., and Ritter, P. (2017). The dynamics of resting fluctuations in the brain: metastability and its dynamical cortical core. Scientific Reports 7, 3095.

di Santo, S., Villegas, P., Burioni, R., and Muñoz, M.A. (2017). Simple unified view of branching process statistics: Random walks in balanced logarithmic potentials. Phys Rev E 95, 032115.

Dickey, A.S., Suminski, A., Amit, Y., and Hatsopoulos, N.G. (2009). Single-unit stability using chronically implanted multielectrode arrays. J Neurophysiol 102, 1331–1339.

Donoghue, J.P., Sanes, J.N., Hatsopoulos, N.G., and Gaál, G.n. (1998). Neural discharge and local field potential oscillations in primate motor cortex during voluntary movements. J Neurophysiol 79, 159–173.

Fox, M.D., and Raichle, M.E. (2007). Spontaneous fluctuations in brain activity observed with functional magnetic resonance imaging. Nat Rev Neurosci 8, 700–711.

Fraiman, D., and Chialvo, D.R. (2012). What kind of noise is brain noise: anomalous scaling behavior of the resting brain activity fluctuations. Front Physiol 3, 307.

Fraser, G.W., and Schwartz, A.B. (2011). Recording from the same neurons chronically in motor cortex. J Neurophysiol 107, 1970–1978.

Gautam, H., Hoang, T.T., McClanahan, K., Grady, S.K., and Shew, W.L. (2015). Maximizing sensory dynamic range by tuning the cortical state to criticality. PLoS Comput Biol 11, e1004576.

Hansen, E.C.A., Battaglia, D., Spiegler, A., Deco, G., and Jirsa, V.K. (2015). Functional connectivity dynamics: Modeling the switching behavior of the resting state Neuroimage 105, 525–535.

Harris, T.E. (1963). The Theory of Branching Processes (Berlin: Springer-Verlag).

Honey, C.J., Kotter, R., Breakspear, M., and Sporns, O. (2007). Network structure of cerebral cortex shapes functional connectivity on multiple time scales. Proc Natl Acad Sci USA 104, 10240–10245.

Hutchison, R.M., Womelsdorf, T., Allen, E.A., Bandettini, P.A., Calhoun, V.D., Corbetta, M., Della Penna, S., Duyn, J.H., Glover, G.H., Gonzalez-Castillo, J., et al. (2013). Dynamic functional connectivity: Promise, issues, and interpretations Neuroimage 80, 360–378.

Jackson, A., and Fetz, E.E. (2007). Compact movable microwire array for long-term chronic unit recording in cerebral cortex of primates. J Neurophysiol 98, 3109–3118.

Katzner, S., Nauhaus, I., Benucci, A., Bonin, V., Ringach, D.L., and Carandini, M. (2009). Local origin of field potentials in visual cortex Neuron 61, 35–41.

Kinouchi, O., and Copelli, M. (2006). Optimal dynamical range of excitable networks at criticality. Nat Phys 2 348–351.

Kirst, C., Timme, M., and Battaglia, D. (2016). Dynamic information routing in complex networks. Nature communications 7, 11061.

Klaus, A., Yu, S., and Plenz, D. (2011). Statistical analyses support power law distributions found in neuronal avalanches. PloS one 6, e19779.

Laurson, L., Illa, X., Santucci, S., Tore Tallakstad, K., Måløy, K.J., and Alava, M.J. (2013). Evolution of the average avalanche shape with the universality class. Nature communications 4, 2927.

Levina, A., and Priesemann, V. (2017). Subsampling scaling. Nature communications 8, 15140.

Li, M., Liu, F., Jiang, H., Lee, T.S., and Tang, S. (2017). Long-term two-photon imaging in awake macaque monkey Neuron 93, 1049–1057.e1043.

Liu, X., and Duyn, J.H. (2013). Time-varying functional network information extracted from brief instances of spontaneous brain activity. Proc Natl Acad Sci U S A 110, 4392–4397.

Lutcke, H., Margolis, D.J., and Helmchen, F. (2013). Steady or changing? Long-term monitoring of neuronal population activity. Trends Neurosci 36, 375–384.

Ma, Z., Turrigiano, G.G., Wessel, R., and Hengen, K.B. (2019). Cortical circuit dynamics are homeostatically tuned to criticality in vivo Neuron 104, 655–664.

Mehring, C., Rickert, J., Vaadia, E., Cardosa, d.O., Aertsen, A., and Rotter, S. (2003). Inference of hand movements from local field potentials in monkey motor cortex. Nat Neurosci 6, 1253–1254.

Meisel, C., Olbrich, E., Shriki, O., and Achermann, P. (2013). Fading signatures of critical brain dynamics during sustained wakefulness in humans. J Neurosci 33, 17363–17372.

Miller, S.R., Yu, S., and Plenz, D. (2019). The scale-invariant, temporal profile of neuronal avalanches in relation to cortical γ–oscillations. Scientific Reports 9, 16403.

Nicolelis, M.A., Dimitrov, D., Carmena, J.M., Crist, R., Lehew, G., Kralik, J.D., and Wise, S.P. (2003). Chronic, multisite, multielectrode recordings in macaque monkeys. Proc Natl Acad Sci USA 100, 11041–11046.

Onnela, J.P., Saramaki, J., Hyvonen, J., Szabo, G., Lazer, D., Kaski, K., Kertesz, J., and Barabási, A.L. (2007). Structure and tie strengths in mobile communication networks. Proc Natl Acad Sci USA 104, 7332–7336.

Pajevic, S., and Plenz, D. (2009). Efficient network reconstruction from dynamical cascades identifies small-world topology from neuronal avalanches. PLoS Comp Biol 5, e1000271.

Pajevic, S., and Plenz, D. (2012). The organization of strong links in complex networks NatPh 8, 429–436.

Pasquale, V., Massobrio, P., Bologna, L.L., Chiappalone, M., and Martinoia, S. (2008). Self-organization and neuronal avalanches in networks of dissociated cortical neurons Neurosci 153, 1354–1369

Petermann, T., Thiagarajan, T., Lebedev, M.A., Nicolelis, M.A., Chialvo, D.R., and Plenz, D. (2009). Spontaneous cortical activity in awake monkeys composed of neuronal avalanches. Proc Natl Acad Sci USA 106, 15921–15926.

Plenz, D. (2012). Neuronal avalanches and coherence potentials. European Physical Journal Special Topics 205, 259–301.

Ponce-Alvarez, A., Jouary, A., Privat, M., Deco, G., and Sumbre, G. (2018). Whole brain neuronal activtity displays crackling noise dynamics Neuron 100, 1446–1459.e1446.

Radicchi, F., Castellano, C., Cecconi, F., Loreto, V., and Parisi, D. (2004). Defining and identifying communities in networks. Proc Natl Acad Sci USA 101, 2658–2663.

Raichle, M.E. (2015). The brain’s default mode network. Annu Rev Neurosci 38, 433–447.

Rasch, M.J., Gretton, A., Murayama, Y., Maass, W., and Logothetis, N.K. (2008). Inferring spike trains from local field potentials. J Neurophysiol 99, 1461–1476.

Ribeiro, T.L., Copelli, M., Caixeta, F., Belchior, H., Chialvo, D.R., Nicolelis, M.A., and Ribeiro, S. (2010). Spike avalanches exhibit universal dynamics across the sleep-wake cycle. PloS one 5, e14129.

Ribeiro, T.L., Ribeiro, S., Belchior, H., Caixeta, F., and Copelli, M. (2014). Undersampled critical branching processes on small-world and random networks fail to reproduce the statistics of spike avalanches. PloS one 9, e94992.

Sethna, J.P., Dahmen, K.A., and Myers, C.R. (2001). Crackling noise Nature 410, 242–250.

Shew, W.L., Yang, H., Petermann, T., Roy, R., and Plenz, D. (2009). Neuronal avalanches imply maximum dynamic range in cortical networks at criticality. J Neurosci 29, 15595–15600.

Shew, W.L., Yang, H., Yu, S., Roy, R., and Plenz, D. (2011). Information capacity is maximized in balanced cortical networks with neuronal avalanches. J Neurosci 5, 55–63.

Shriki, O., Alstott, J., Carver, F., Holroyd, T., Henson, R.N., Smith, M.L., Coppola, R., Bullmore, E., and Plenz, D. (2013). Neuronal avalanches in the resting MEG of the human brain. J Neurosci 33, 7079–7090.

Stewart, C.V., and Plenz, D. (2007). Homeostasis of neuronal avalanches during postnatal cortex development in vitro. J Neurosci Meth 169, 405–416.

Tagliazucchi, E., Balenzuela, P., Fraiman, D., and Chialvo, D.R. (2012). Criticality in large-scale brain fMRI dynamics unveiled by a novel point process analysis. Front Physiol 3.

Tetzlaff, C., Okujeni, S., Egert, U., Wörgötter, F., and Butz, M. (2010). Self-organized criticality in developing neuronal networks. PLoS Comput Biol 6, e1001013.

Tolias, A.S., Ecker, A.S., Siapas, A.G., Hoenselaar, A., Keliris, G.A., and Logothetis, N.K. (2007). Recording chronically from the same neurons in awake, behaving primates. J Neurophysiol 98, 3780–3790.

Villegas, P., Di Santo, S., Burioni, R., and Muñoz, M.A. (2019). Time-series thresholding and the definition of avalanche size. Phys Rev E 100.

Watts, D.J., and Strogatz, S.H. (1998). Collective dynamics of ‘small-world’ networks Nature 393, 440–442.

Yang, H., Shew, W.L., Roy, R., and Plenz, D. (2012). Maximal variability of phase synchrony in cortical networks with neuronal avalanches. J Neurosci 32, 1061–1072.

Yu, S., Klaus, A., Yang, H., and Plenz, D. (2014). Scale-invariant neuronal avalanche dynamics and the cut-off in size distributions. PLoS One 9, e99761.

Yu, S., Ribeiro, T.L., Meisel, C., Chou, S., Mitz, A., Saunders, R., and Plenz, D. (2017). Maintained avalanche dynamics during task-induced changes of neuronal activity in nonhuman primates eLife 6, e27119.

Yu, S., Yang, H., Nakahara, H., Santos, G.S., Nikolic, D., and Plenz, D. (2011). Higher-order interactions characterized in cortical activity. J Neurosci 31, 17514–17526.

Zalesky, A., Fornito, A., Cocchi, L., Gollo, L.L., and Breakspear, M. (2014). Time-resolved resting-state brain networks. Proc Natl Acad Sci U S A 111, 10341.

